# Symmetry and force response of cohesin loop extrusion are determined by diffusion of its motor and anchor domains

**DOI:** 10.64898/2026.06.08.730904

**Authors:** Georgii Pobegalov, Runze Sha, Oleg Dobrokhotov, Torahiko Higashi, Chris A. Brackley, Frank Uhlmann, Maxim Molodtsov

## Abstract

Cohesin, a Structural Maintenance of Chromosomes (SMC) complex is thought to organize genomes by generating DNA loops, yet the molecular basis of loop extrusion remains incompletely understood. Here, we show that a single *S. pombe* cohesin complex extrudes DNA loops both symmetrically or asymmetrically, depending on the external force applied to DNA and that the mode and speed are controlled by stochastic switching of the cohesin motor between driving and diffusive states. We directly measured the weak forces generated during cohesin-mediated loop extrusion while simultaneously visualizing DNA loops and found that forces as low as 0.05 pN pull DNA out of cohesin loop without disrupting cohesin–DNA interaction. We further identified the Scc3 subunit as a diffusive DNA anchor required for loop extrusion and showed that its physical separation from the motor domain by unstructured regions of Scc1 promotes loop-extrusion initiation. Together, our results support a model in which cohesin loop extrusion is limited and tuned by diffusive motion of both anchor and motor modules, providing a mechanistic framework for how weak and diffusive cohesin–DNA interactions could modulate cohesin function in chromatin organization.

## Introduction

Cohesin is a eukaryotic protein complex from the Structural Maintenance of Chromosome (SMC) family that plays crucial roles in sister chromatid cohesion, genome organisation, gene expression and DNA repair [1–5]. Cohesin function in gene expression and DNA organization are thought to be facilitated by its ability to move on DNA and extrude DNA loops using the energy of ATP hydrolysis [6, 7]. However, molecular mechanisms used by cohesin to extrude DNA loops remain incompletely understood [8, 9].

The core of cohesin consists of two ∼ 50 nm long Smc1 and Smc3 coiled-coil subunits that dimerize at the globular hinge domain and transiently interact via ATPase head domains. The head domains are also connected by a partially unstructured Scc1 kleisin subunit with all three proteins forming a tripartite ring-like structure (Fig. 1a). The fourth subunit, Scc3, binds Scc1 and provides additional interfaces for interactions with both DNA and the cohesin loader Scc2 [4, 10, 11].

**Figure 1.**
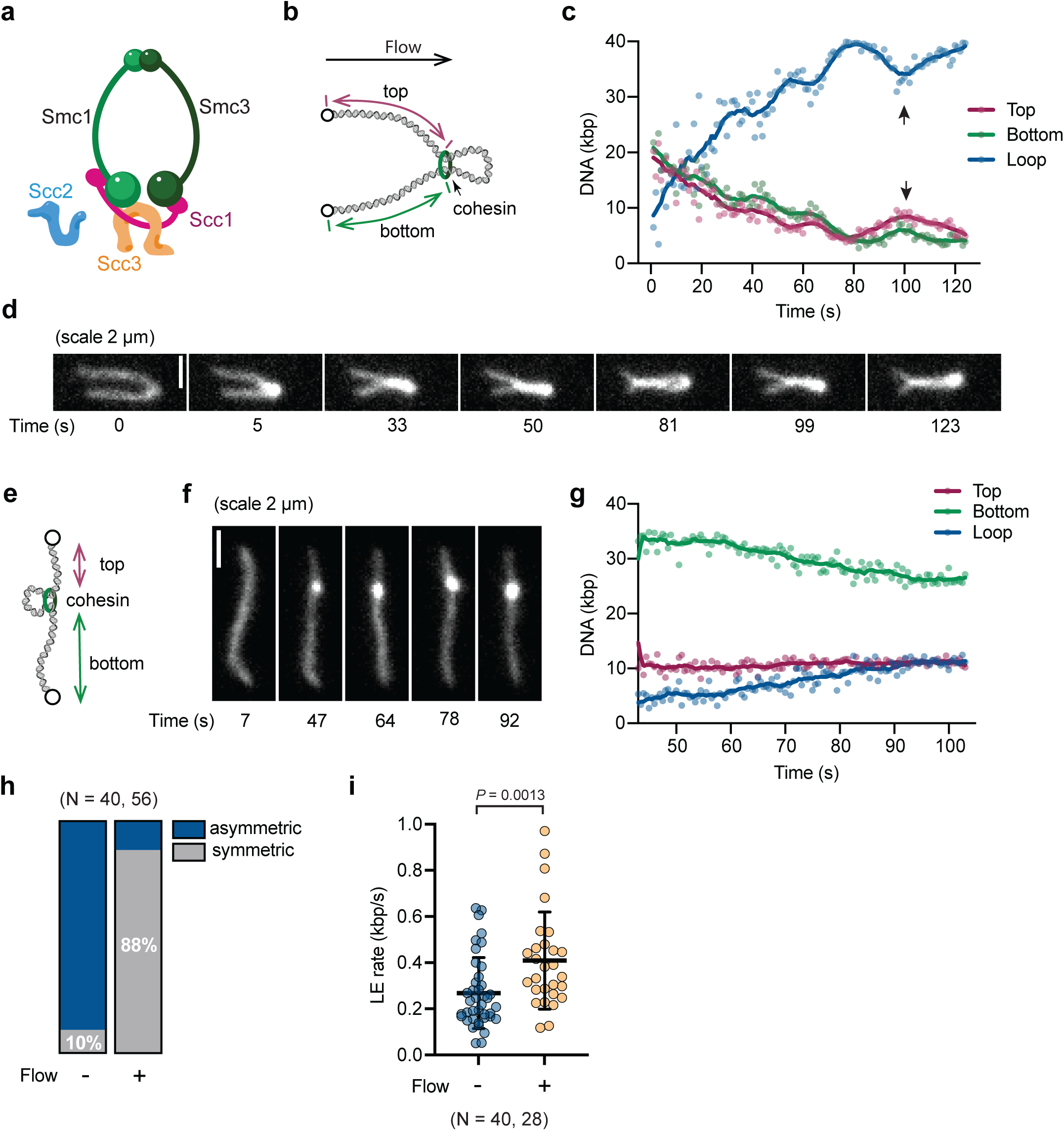
Single cohesin extrudes DNA loops symmetrically in the presence of flow and asymmetrically without it. (a) Schematics of the cohesin complex. Individual subunits are indicated using budding yeast nomenclature; (b) Schematics of the DNA in the loop extrusion assay with the flow. Cohesin is shown at the loop base (green), and loop growth is quantified by tracking the shortening of the two non-looped arms shown as “top” and “bottom”; (c) Time trace quantification showing the lengths of the top (magenta) and bottom (green) DNA arms during loop extrusion under flow, illustrating near-symmetric incorporation of DNA into the loop. Arrowheads point to an event where cohesin “slides back” reducing the size of the loop; (d) Representative snapshots from a movie showing symmetric loop elongation corresponding to c; (e) Schematic of experiment without the flow. Two DNA arms outside the loop defined as the top and bottom segments; (f) Representative fluorescence images of a loop-extrusion without the flow showing progressive loop growth over time seen as increased lump on DNA; Scale bar, 2 μm; (g) Representative arm and loop length traces. In the absence of flow, one DNA arm shortens whereas the other remains nearly constant; (h) Quantification of loop-extrusion symmetry in the presence and absence of flow. (i) Distribution of loop-extrusion (LE) rates measured in the presence and absence of flow. Horizontal lines represent the mean and standard deviation values.

Cohesin can efficiently extrude DNA loops on naked DNA in vitro and the underlying molecular mechanism of loop extrusion has been subject of intense experimental and theoretical study. Although different models have been proposed, they all agree that the main driver of cohesin movement is ATP-dependent changes of the head domains coupled with dynamics of SMC coiled-coils, which result in one-directional translocation along the DNA [12–14]. However, to both generate and maintain a loop an additional interaction site with the second DNA strand is required alongside the translocating unit. Thus, existing models predict that a single SMC can only extrude loops asymmetrically as the motor domain can only actively move on one DNA strand at a time. Because one-sided loop extrusion is often not sufficient by itself to explain the full range of chromosome-organization features [15], alternative scenarios were proposed and received experimental support. First, it could be that cohesin can switch DNA strands during loop extrusion resulting in effective symmetric movement [13, 16, 17]. Alternatively, cohesin may extrude DNA loops as a dimer motoring on both DNA strands at the same time [6, 18], consistent with observations made in living cells [19]. Since different studies suggest that loop extrusion may be mediated either by monomeric cohesin or by cohesin dimers, the relationship between cohesin’s oligomeric state and its loop extrusion activity in vitro remains unresolved.

In contrast to loop elongation, much less is known about how cohesin complexes initiate DNA loops in the first place. Given the size of the cohesin complex is approximately equal to the DNA persistence length, it was proposed that cohesin may need to actively bend DNA using ATP dependent head-head engagement for efficient initiation [20]. Alternatively, loops formed spontaneously due to random thermal fluctuations of DNA shape could serve as seeds for loop extrusion trapped by cohesin at the initial stages [21]. However, how cohesin can capture and rectify these thermally formed DNA loops is not well understood.

Cohesin as well as other SMCs are also known to be very weak motors that stall at forces ∼ 0.1 pN [13, 22, 23]. In addition, loop extrusion can sometimes be interspersed with loop diffusion and loop slipping events suggesting that diffusive cohesin/DNA interactions are part of the cohesin activity [13, 17]. Cohesin diffusion may significantly facilitate generation of DNA loops even without motoring activity [24, 25]. However, where such diffusive interactions are located within the cohesin–DNA complex and how they control loop initiation, elongation and determine the symmetry, processivity, and force sensitivity of loop extrusion is still poorly understood.

## Results

### Single cohesin can drive both symmetric and asymmetric loop extrusion

To visualize loop extrusion by yeast cohesin we used an assay in which λ-DNA (48.5 kbp long) is tethered at two ends to the passivated glass surface of a microfluidic flow cell (Fig. 1b). DNA was stained with Sytox Orange and imaged using highly inclined and laminated optical sheet (HILO) microscopy in the presence of a mild flow. In the presence of cohesin, Scc2 loader and ATP we observed formation and processive growth of individual DNA loops (Fig. 1c,d, Video S1). We quantified loop growth by measuring the length of two DNA segments from each tethering point to the base of the loop (Fig. 1b), corresponding to remaining DNA outside of the loop. As the loop grows the length of these two segments decrease. Similar to what has been reported for human cohesin [7], yeast cohesin effectively extruded DNA loops very symmetrically (Fig. 1c,d). Out of n=56 traces, 49 (88%) showed phases of quick symmetric loop extrusion with speeds reaching hundreds of base pairs per second, which was frequently followed by slipping and diffusing events when the base of the loop approached DNA tethering points.

Human cohesin has been shown to act during loop extrusion as a monomer or as a dimer in different reports [6, 7], and in yeast the number of cohesin complexes required for loop extrusion has not yet been determined. To test whether symmetric loop extrusion was the result of more than one cohesin complex acting on two DNA strands we labelled cohesin with Alexa-647 dye at the C-terminus of Scc1 (Fig. S1a,b). We used two-colour imaging to simultaneously visualise both DNA and cohesin and quantified cohesin stoichiometry by photobleaching Alexa-647. Labelled cohesin was visible at the bases of the loops in 81% cases and in 59% cases (n=81) the fluorescent intensity corresponded to a single cohesin (Fig. S1d-f). Given that cohesin labelling efficiency was 86%, our results strongly suggest that a single yeast cohesin complex can efficiently extrude DNA loops *in vitro*.

Symmetric loop extrusion could be a result of switching between different DNA strands, where cohesin moves on one strand and then switches and moves on the other, which has been proposed theoretically and observed in experiments without flow [13, 17]. However previously observed switching times were on the order of ∼ 45 s on average [17], which could not explain fast symmetric loop extrusion events observed in our experiments: in the presence of flow, DNA is incorporated into the loop from both sides in under 1 s resolution (Fig. 1c). Therefore, we wondered whether externally applied force by the flow may facilitate faster and more symmetric loop extrusion. To test this, we compared our results in the presence of flow to the identical conditions but without the flow. Indeed, similarly to human cohesin, yeast cohesin showed predominantly asymmetric loop extrusion without the flow (Fig. 1e-h, Video S2). Interestingly the speed of loop extrusion was also significantly slower (Fig. 1i). Thus, external flow that stretches DNA also increases the speed of loop extrusion and makes the process symmetric.

### Model with diffusive motor and anchor domains accounts for loop extrusion with and without flow

Next, we asked whether the simplest possible model of cohesin with one motor domain and one diffusive anchor domain could explain the differences in the loop extrusion in the presence and absence of flow. To this end, we used coarse-grained molecular dynamics simulations and represented cohesin as two units that can move along DNA, which was represented as a chain of beads. Cohesin motoring and diffusive units were connected by a finite extensible nonlinear elastic spring that limited extent of its stretching to 50 nm – approximately the size of the cohesin complex. With this model, we investigated how movement of cohesin in the presence and absence of flow depends on the diffusion of the anchor unit and the state of the motoring unit (Fig. 2a). Motoring was implemented by having the motor cohesin unit attached to a given DNA bead and then removing and replacing this attachment with one connecting to the next DNA bead, repeating this process at a constant rate. In contrast, diffusive interaction was specified such that thermal fluctuations could drive the diffusing unit along the DNA molecule with the attraction ensuring it remained in contact (see Methods for details).

**Figure 2.**
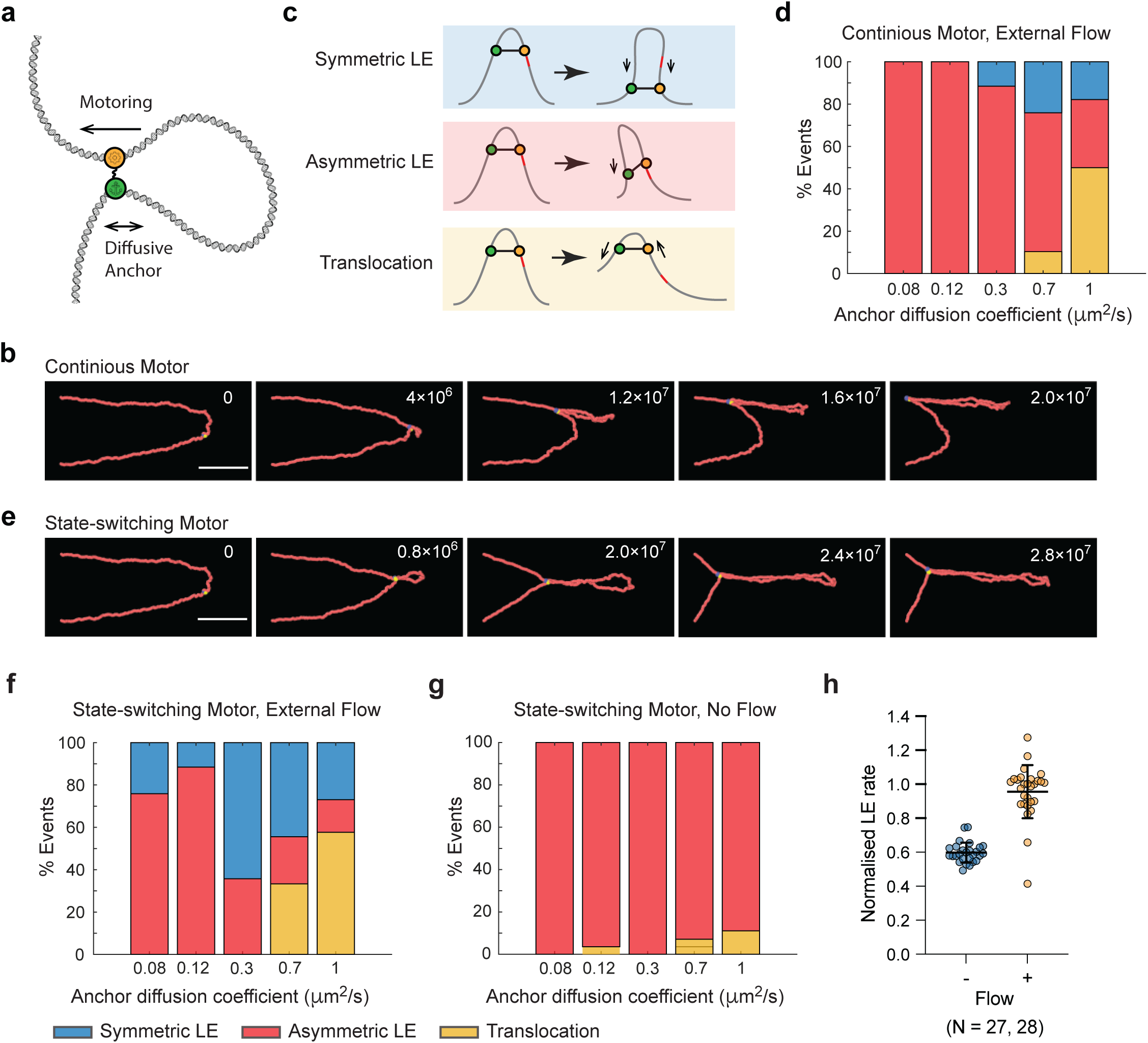
Minimal model explains symmetric and asymmetric loop extrusion. (a) Schematics of the model in which cohesin is represented by two units - a motor and an anchor; (b) Snapshots from the simulation with slowly diffusing anchor and continuously moving cohesin motor units. Blue and yellow beads represent motor and anchor cohesin units respectively. Scale bar is 1 µm and numbers show iteration number of the simulation. Size of the beads was artificially increased for visualization purposes. Diffusion coefficient of the anchor is 0.3 µm^2^/s; (c) Schematics of the symmetric, asymmetric loop extrusion and translocation events. Red segment marks discrete position on DNA to assist visualization. Arrows show net direction of the motor and anchor movement; (d) Quantification of events in the presence of flow for different diffusion coefficients of anchor in the model in which motor unit continuously moves until the DNA tether point is reached; Blue, red and yellow stand for symmetric, asymmetric and translocation events as shown in c. Each column is based on 30 simulations; (e) Snapshots from the simulation in which the motor stochastically switches between motoring and diffusive states. Other parameters and notations as in (b); (f) Quantification of events in the presence of flow for a model in which cohesin motoring unit switches between motoring and diffusive states; (g) Quantification of events without flow for the model in which cohesin motoring unit switches between motoring and diffusive states; (h) Comparison of the loop extrusion rates with and without the flow for diffusion coefficient of 0.3 µm^2^/s. Loop extrusion rate is normalized to the mean rate in the presence of flow. Horizontal lines represent the mean and standard deviation values.

As expected, a slowly diffusing anchor resulted in mostly asymmetric loop extrusion when the movement of the motor was continuous (Fig. 2b, Video S3). To quantify the asymmetry of the cohesin loop extrusion we introduced a directionality parameter 𝜉 as the ratio between the distance travelled by each unit of cohesin towards DNA tether points. Thus, 𝜉 =1 corresponds to ideally symmetric loop extrusion (both anchor and motor travel the same distance in opposite directions towards their respective DNA tether points increasing the loop size), 0 for asymmetric (the anchor is completely immobile) and -1 for translocation (both units move towards the same DNA anchor point with the same speed).

Next, we varied the strength of the interaction between the anchor and DNA, which controls the anchor diffusion coefficient, and classified all outcomes as either symmetric, asymmetric loop extrusion or translocation (Fig. 2c, for details see methods). For each value of the attraction strength, we simulated a minimum of 30 trajectories, and to simulate the flow, drag force was applied to all DNA and cohesin beads. Our simulations showed that in the presence of flow, outcomes of the simulation depended on the diffusion coefficient of the anchor. As the diffusion coefficient increased and it could move more freely on DNA, and this was associated with were fewer asymmetric loop extrusion events and more events of the other types (Fig. 2d). For a highly diffusive anchor we observed both translocation and symmetric loop extrusion events (Video S4 and S5). Although translocation events were unexpected, they were indeed occasionally observed experimentally [7, 26]. Symmetric loop extrusion events also occurred occasionally but represented less than 20% of all events and thus no and consistent symmetric loop extrusion was observed.

Since our experiments suggested that the motoring cohesin unit can switch stochastically between motoring and diffusing states, we next tested the consequences of such switching. To this end, we simulated a scenario in which the cohesin motor domain instead of moving continuously in one direction switched several times during the simulation into a diffusive mode and then back into the driving mode (on average up to six times before reaching the DNA tether point). The other cohesin unit was always in the diffusing state. These simulations resulted in a much larger proportion of symmetric loops and significantly more stable symmetric loop extrusion in the presence of flow (Fig. 2e, Video S6). This is because when the motor switches into a diffusive state, DNA strand with cohesin closer to the tether point experiences stronger tension due to external flow, which reintroduces the symmetry by aligning the loop along the flow. When the motoring unit switches back from the diffusive to the motoring state, motoring along the symmetric configuration has higher chance of biasing the diffusion of the anchor subunit towards the direction of the motor movement resulting in symmetric loop extrusion similar to that observed experimentally (Fig. 1d). Our simulations showed that there is an optimal diffusion coefficient of the anchor at which ∼70% of simulations resulted in symmetric loop extrusion whereas more diffusive anchor leads to more translocation events and more stable anchor leads to more asymmetric loop extrusion (Fig. 2f).

To further validate our model, we also performed simulations where the motoring unit switched states, but no external flow was present. None of the simulations in this case resulted in symmetric loop extrusion. This is expected because without the symmetry imposed on DNA by the flow the anchor does not have any preferred direction of diffusion regardless of the movement of the motoring unit (Fig. 2g). The simulations also showed that loop extrusion speed in this case was smaller than in the presence of flow (Fig. 2h) in agreement with our experiments (Fig. 1i). Thus, a simple model representing cohesin as the motor and diffusive anchor subunits connected by flexible linkage fully explains loop extrusion *in vitro* both with and without the flow.

### Diffusive cohesin/DNA interaction limits its ability to generate force

The diffusive nature of the cohesin/DNA interaction at both anchor and diffusive motor states should affect not only loop extrusion symmetry, but also behaviour of cohesin under external force. To probe it more directly, we applied controlled tension to one end of the DNA using an optical trap while visualizing DNA loops using HILO microscopy (Fig. 3a). To achieve this, we tethered one end of DNA to the surface of the flow cell and the other end to the 1-micron sized bead. The bead was held in an optical trap allowing for dynamic changes of the DNA end-to-end distance, simultaneous measurement of the DNA tension and its visualization.

**Figure 3.**
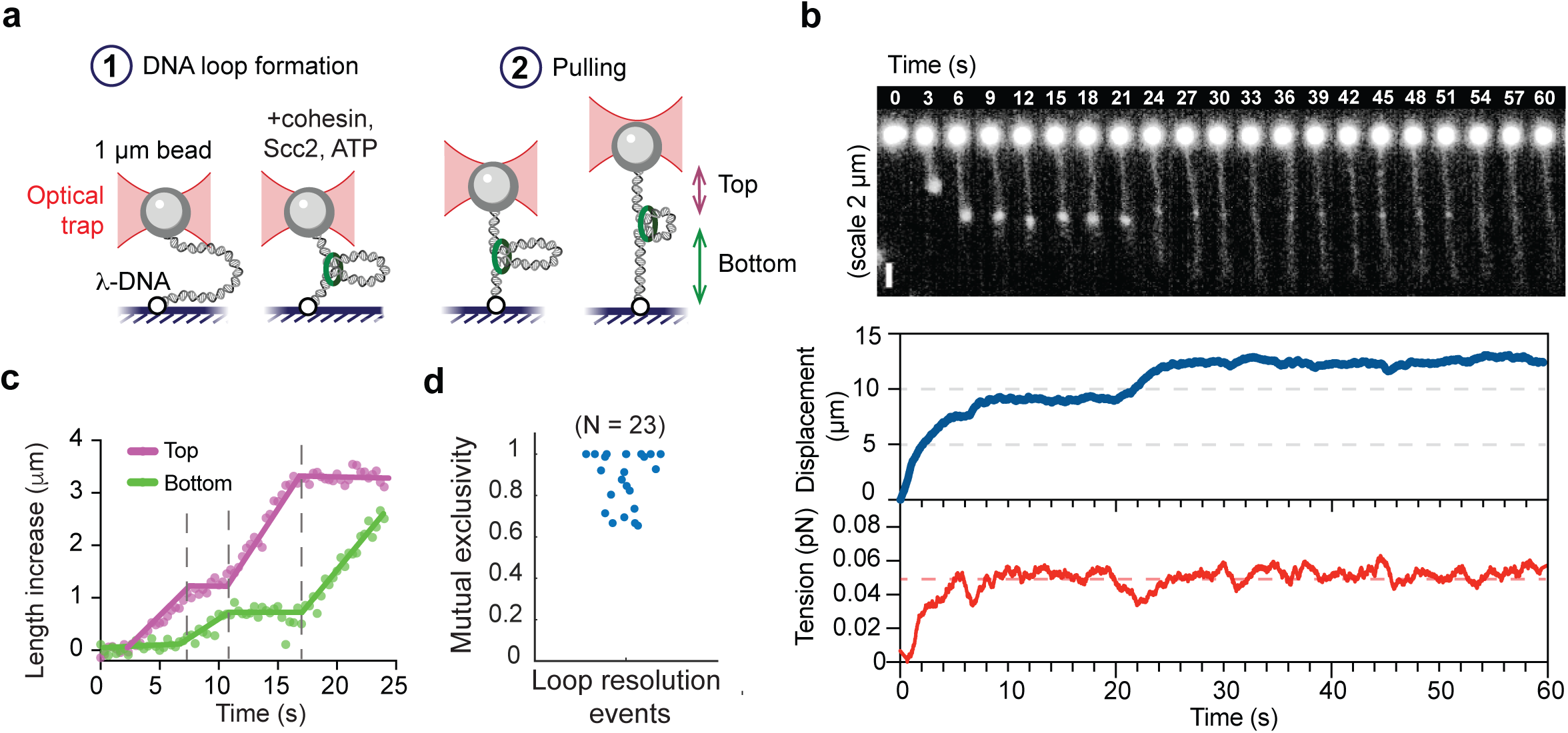
Small external force releases DNA asymmetrically from cohesin generated loops. (a) Schematics of the assay. In step 1 DNA loop is formed by bringing DNA ends closer together. In the step 2, DNA is stretched by moving the bead. The “top” and “bottom” DNA segments flanking the loop are defined by the tether geometry; (b) pulling on a DNA loop at a constant tension of 0.05 pN. Top: Representative fluorescence time series; middle: distance between the DNA ends, scale bar, 2 µm; bottom: the corresponding DNA tension; (c) Representative example of the loop release dynamics. Dots are experimental points and lines are trendlines. Dashed lines denote points at which DNA release is switched between the top and bottom arms; (d) Mutual exclusivity index shows fraction of time during which DNA was pulled from one side of the loop only. Each datapoint corresponds to one independent experiment.

To form a loop, we incubated DNA in a relaxed state in the presence of cohesin, loader and ATP by bringing the optical trapped bead with one end of the DNA to the anchor point of the other end. After DNA loop formation was confirmed by imaging, we pulled on DNA by moving the bead away parallel to the coverslip and visualized DNA loop dynamics (Fig. 3b, Fig. S1g, Video S7). In all cases, after we applied force, loops were gradually resolved as DNA was pulled out of the loop and no loop extrusion against external tension was observed (n = 23). The lowest force we could apply in this setup was ∼ 0.05 pN, which was limited by the Brownian forces acting on the bead. Even at this low force we could only see loop dissolution events, which shows that cohesin cannot extrude loops against forces of ∼ 0.05 pN and that this small external force is enough to promote DNA sliding out of the loop formed by cohesin.

Interestingly, in majority of cases loop resolution was asymmetric. Under applied tension DNA was released predominantly from one side of the loop, while the DNA length on other side stayed fixed. Occasionally the two sides switched, and the DNA was released from the other side of the loop (Fig. 3c). Some traces demonstrated more than one switch back and forth but in 90% cases (40 switches out of 44), DNA switched fully from being released at one side and locked on the other side to completely the opposite – locked at the first side and released in the other. To quantify this, we calculated mutual exclusivity index as a fraction of time for each trace that DNA is being pulled only from one side (as opposed to being pulled from both sides at the same time or not being pulled at all). This index showed that across all traces more than 90% of the time DNA was pulled only from one side of the loop (Fig. 3d).

These results are accounted for by our model in which cohesin contains a diffusive anchor and a motor domain that stochastically switches between motoring and diffusive states. Under applied external force, the motor and anchor are subject to the same DNA tension. As a result, DNA will preferentially slide out of the loop through whichever interface presents the lower effective resistance (i.e., lower friction) to sliding. When cohesin is in the motoring state, it is strongly engaged with DNA and therefore must provide a higher-resistance contact with DNA than the diffusive anchor. Thus, DNA is expected to slip out primarily at the anchor, producing asymmetric loop shrinkage. When the motor stochastically switches into a diffusive state, both interfaces (diffusive motor and diffusive anchor) can, in principle, permit sliding. In that regime, the exit side is determined by their relative frictions: DNA will slide out through the interface with the lower friction. The observed switching events therefore imply that the motor, when in its diffusive state, offers less resistance to DNA sliding than the anchor, causing the preferred exit side to flip accordingly. Thus, weak-force, one-sided release of DNA from cohesin loops strongly supports the idea that loop extrusion is constrained by two frictional DNA-binding interfaces.

### The diffusive anchor is cohesin Scc3 subunit

While cohesin motoring is presumably driven by the action of Smc subunits, it is much less well understood which part of the cohesin complex corresponds to the diffusive anchor. Previously, we and others showed that removing DNA binding site on the Scc3 subunit by replacing lysins with glutamates abrogates ability of cohesin to initiate DNA loops [27, 28]. Since anchor/DNA interaction are required for loop extrusion, we reasoned this DNA binding site might be good candidate for the diffusive anchor.

The overall organization of the Scc3 subunit in the cohesin complex also matches well with our model in which motor and anchor domains are connected by a flexible linkage (Fig. 4a). Indeed, Scc3 is connected to Smc1 and Smc3 subunits via flexible unstructured domains of Scc1 that can stretch to over 50 nm [13, 29]. In combination with the observation that Scc3 DNA binding is required to start loop extrusion, this suggests two possible scenarios for the loop extrusion initiation. In the first scenario cohesin passively rectifies a small DNA loop generated by random thermal fluctuations (Fig. 4b, Scenario 1). In the second scenario, motor and anchor domains bind linear stretch of DNA and movement of the motor generates initial DNA loop (Fig. 4b, Scenario 2). Both scenarios predict that efficiency of loop initiation must depend on the length of the flexible linker between the motor and anchor domains. Indeed, short linker requires generation of a smaller and steeper initial DNA loop by random thermal fluctuations for cohesin to start loop extrusion. At the same time longer linker would allow loop extrusion to initiate with less DNA bending. Stronger DNA bending requires more highly energetic thermal fluctuations, which are less likely. Thus, we would expect cohesin with longer linker between the motor and anchor domains to initiate loop extrusion more effectively.

**Figure 4.**
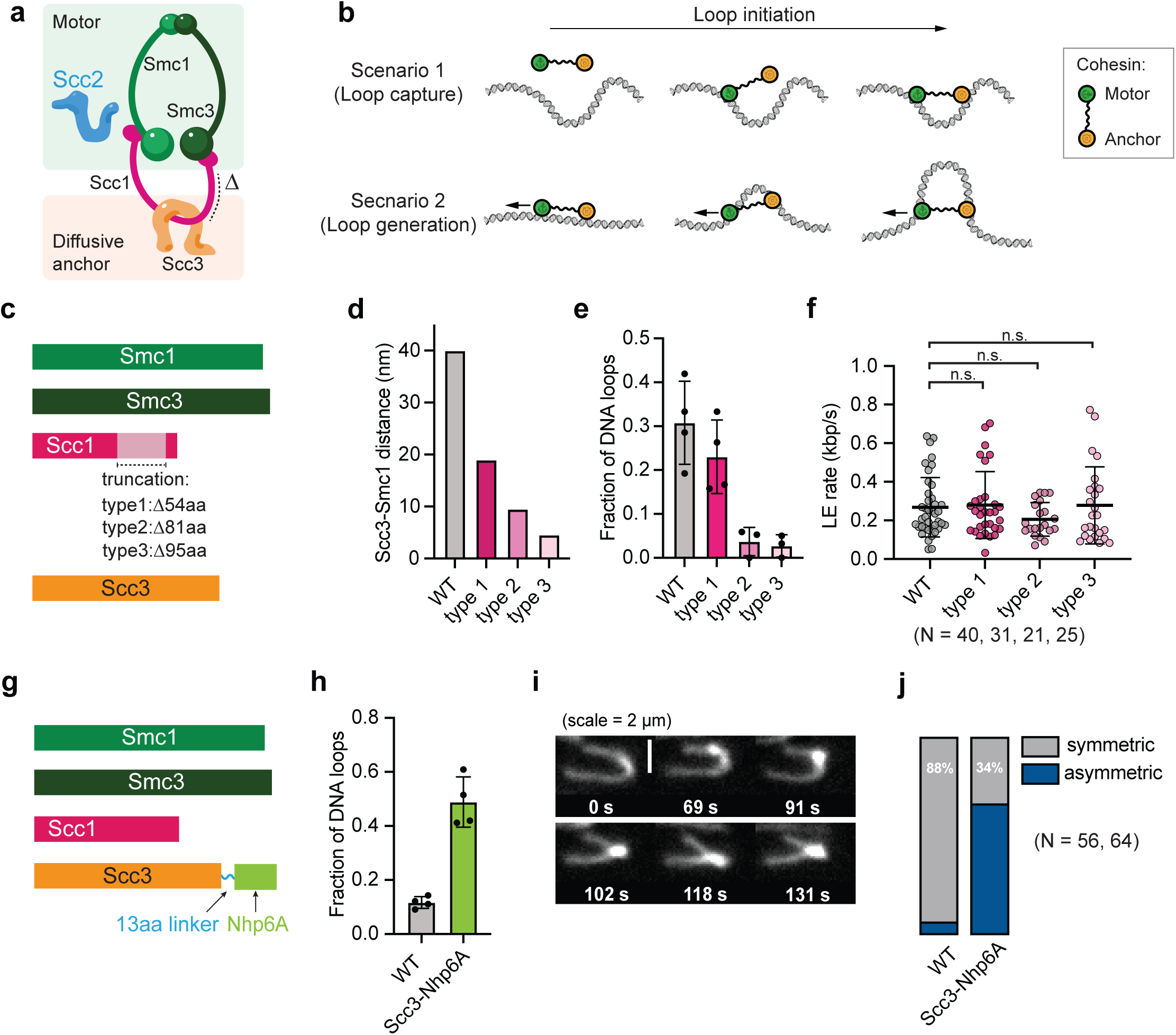
Scc1 and Scc3 subunits determine anchor position and control the symmetry and initiation efficiency of loop extrusion. (a) Schematics of the motor and anchor subunits. Δ indicates position of Scc1 truncation site used in (c); (b) Two proposed scenarios for loop initiation. In scenario 1 cohesin captures DNA loop already pre-formed by random thermal fluctuations. In scenario 2, cohesin binds straight DNA and motoring subunit moves away from the anchor (shown with an arrow), which facilitates DNA loop formation; (c) Schematics of cohesin construct with shortened Scc1 at C-terminus; (d) Calculated length of the C-terminal fragment of Scc1 connecting Smc1 and Scc3; (e) Loop-extrusion initiation efficiencies for cohesin variants with progressively shortened Scc1 C-termini. All complexes were used at the same concentrations. Efficiencies are measured as a fraction of DNA molecules showing loops over 10 minutes observation period (n = 115, 74, 78 and 71 DNA molecules for WT, type1, type2 and type3 cohesin respectively). Lines show standard deviations; (f) Loop extrusion rate distributions for wild type cohesin and Scc1-shortened variants. Horizontal lines represent the mean and standard deviation values; (g) Schematics of the chimeric construct with Nhp6 fused to C-terminus of Scc3; (h) Loop extrusion efficiency comparing wild type and Nhp6A-chimeric cohesin (n = 127 and 121 DNA molecules for WT and Nhp6A-Scc3 cohesin respectively). Lines show standard deviations; (i) Representative time series for Nhp6A chimera cohesin showing asymmetric loop extrusion. Scale bar 2 μm; (j) Quantification of loop extrusion symmetry for WT and Nhp6A-Scc3 cohesin in the presence of flow, scored as two-sided (symmetric) versus one-sided (asymmetric) extrusion (n = 56, 64).

To test this idea, we generated a series of mutants with progressively shorter intrinsically disordered C-terminal domains of Scc1 with expected lengths of 19, 9.5 and 4.5 nm in comparison with 45 nm that of wild type (Fig. 4c,d). We verified that these complexes with truncated Scc1 retained the ability to hydrolyse ATP and topologically load onto DNA similar to wild type cohesin (Fig. S2a-c). However, in the loop extrusion assays, our experiments show that shorter Scc1 significantly impair ability of cohesin to start loop extrusion (Fig. 4e). This is consistent with our prediction that the separation between Scc1 and ATPase domains is required for initial capture of thermally formed DNA loops. Interestingly, across all types with shortened Scc1, those molecules that did successfully initiate loops could elongate them with rate indistinguishable from wild type complexes showing that once the loop is formed, demonstrating that the distance between the anchor and the motor does not play significant role in loop elongation (Fig. 4f). We also note that while cohesins with shortened disordered domains initiated loop extrusion very inefficiently, these Scc1 variants supported fission yeast cell growth indistinguishable from the wild type (Fig. S2d).

Finally, our model predicts that increasing affinity of the Scc3 DNA binding site to DNA should result in more robust initiation of loop extrusion, but also more asymmetric loop elongation as tighter anchor binding is associated with its slower diffusion. To test this prediction, we have sought to increase DNA binding affinity of Scc3 such that it would nonetheless allow for restricted diffusion along the DNA. To achieve this, we chose budding yeast Nhp6A – an abundant protein that binds DNA non-specifically with nanomolar affinity and can diffuse along DNA [30, 31].

We fused C-terminus of Scc1 via flexible linker to Nhp6A (Fig. 4g) and checked that this chimeric cohesin remained functional and retained the ability to topologically load onto DNA in an ATP-dependent manner (Fig. S3a,b). Also as expected, bulk affinity of this cohesin variant to DNA was increased (Fig. S3c). When tested in loop extrusion assays, the Nhp6A chimera complex showed strongly increased loop initiation with ∼5 times more loops formed in 10 min observation interval (Fig. 4h). Furthermore, our model predicts that adding additional affinity to anchor domain should decrease anchor’s diffusion coefficient and therefore result in more asymmetric loop extrusion events in the presence of flow. In agreement with this prediction, over 60% of Nhp6 chimera molecules (n=64) showed asymmetric loop extrusion comparing to 12% of molecules (n=56) of wild type cohesin (Fig. 4i,j, Video S8).

Taken together, our results support a model in which cohesin loop extrusion is governed by two weak, frictional DNA contacts: a diffusive Scc3 anchor and a motor that stochastically switches between motoring and diffusive states. Our findings show that diffusive cohesin/DNA interaction shapes its extrusion symmetry, force-dependent loop release as well as loop initiation efficiency.

## Discussion

The main conclusion of this work is that diffusive movement of both cohesin anchor and motor domains are an inherent feature of cohesin loop extrusion, which have implications for both loop initiation and elongation. We identified cohesin anchor as residing on the Scc3 subunit, which is mechanically decoupled from the Smc motor domains by the unstructured regions of Scc1. Furthermore, we showed that loop initiation and loop elongation are kinetically distinct steps with different molecular requirements: initiation requires DNA engagement by Scc3 and is constrained by the physical separation between Scc3 and the Smc motor domains, which sets the effective initial capture range for generating DNA loop. In contrast, elongation is independent of the Scc1 length and, as our simulations indicate, proceeds over a range of Scc3–DNA interaction strengths.

A practical implication of this result is that initiation frequency is likely highly context dependent. Any factor that increases the local rate of DNA bending and/or Scc3 affinity to DNA would selectively enhance initiation and leave elongation kinetics unchanged. Furthermore, both Scc1 variants with compromised loop extrusion and loop extrusion deficient cohesin due to mutations in the Scc3 supported wild type level of cell viability (Fig. S2d) [27]. This emphasises the importance of further investigating the possible contribution of *in vitro* observed loop extrusion behaviour to *in vivo* cohesin function.

Our model also resolves the question of why loop extrusion in the presence of flow appear symmetric while asymmetric in its absence. Cohesin is an inherently asymmetric motor due to presence of only one motoring subunit that can move only in one direction at a time. However, when DNA is stretched by the flow, this imposes an external symmetry to the cohesin/DNA system, which makes random movement of the anchor biased in the direction of the motor movement. This also increases the overall speed because the motor and anchor units are moving in the same direction resulting in apparent symmetric loop extrusion with up to double the speed of asymmetrically moving cohesin without the flow. In cells chromatin is subject to continuous thermal motion, but in some cellular contexts external forces may impose sufficient alignment or tension on DNA to bias cohesin toward more symmetric loop extrusion. Most genomic processes including replication and transcription generate DNA tension, which may transiently realign chromatin. Our model shows that such conditions may allow cohesin to switch from asymmetric to symmetric movement. Overall, our work shows that cohesin directionality is determined not simply by its motor but is a net result of the relative friction at two DNA interfaces and the probability of the motor switching between the motoring and diffusive states, which can be easily biased by external force.

In chromatin diffusive states are likely to be modulated by binding to other proteins. For example, although budding yeast lacks CTCF, metazoan genome folding relies heavily on CTCF as a boundary element that stalls cohesin in an orientation-dependent manner [32, 33]. Although cohesin and CTCF have been shown to interact [33, 34], how cohesin movement can be specifically blocked by CTCF remains unknown. Our study suggests that potential solution might be in how CTCF controls diffusive motion of cohesin instead of acting as a simple roadblock. By restricting diffusive motion of cohesin, it might be selectively stopped while interacting with CTCF, but not other proteins that do not affect its diffusion. This view also predicts that the boundary strength should be tuneable by factors that modulate anchor friction, and that the same cohesin complex could display different boundary sensitivities depending on its current motoring vs diffusive state.

Our data are consistent with the idea that the small stalling force of cohesin (∼0.05 pN) likely arises from diffusive interactions between anchoring and motoring subunits with DNA. Thus, even small forces can slide and reposition loop extruding cohesin on DNA. We propose that rather than a weakness, this may be understood as a feature of mechanical compliance. Cohesin remains bound to DNA while allowing DNA to slide through one or the other interface depending on which interface is in a high- vs low-friction state. Most molecular machines including polymerases and helicases generate significantly higher force suggesting that they can easily pull DNA out of (or into) cohesin generated loop [35]. However, while remaining on the DNA, once the external force is removed, cohesin may pull the DNA back into the loop. Thus, the weakness of cohesin as a motor might be a key for maintaining chromatin homeostasis in the face of temporary events requiring forceful DNA rearrangements possibly during replication, repair and especially in mitosis.

Finally, given the large number of cohesins inside the cell [36–38], a strong, highly processive, high-stall-force motor would risk generating overly persistent loops that resist necessary remodelling. By contrast, a weak, diffusion-permissive cohesin is well matched to the need for continuous rearrangement of DNA. Our study identified simple mechanistic rules underlying chromatin plasticity and proposes how it can be potentially controlled by changing diffusive interactions between cohesin and DNA linking molecular mechanism to genome-scale organization. The emerging picture is that chromatin organization is controlled less by the power of an extruder and more by context-dependent regulation of diffusion, friction, and state switching - parameters that are readily tuneable by cohesin binding partners and cell cycle specific regulation.

## Methods

### Protein expression and purification

Recombinant S. pombe cohesin complexes used in this study were expressed, labelled and purified as described previously [27, 35]. Although we used Pombe cohesin everywhere in the text for simplicity we use generic nomenclature, thus Smc1 refers to Psm1, Smc3 to Psm3, Scc1 to Rad21 and Scc3 to Psc3. Briefly cells co-expressing Smc1, Smc3, Scc1, and Scc3 were grown in YP medium containing 2% raffinose to OD600 1.0 at 30 °C, induced with 2% galactose for 2 h, harvested, cryomilled, and lysed in protease-inhibitor-containing buffer with benzonase. After clarification, cohesin was purified by IgG affinity chromatography with on-column PreScission cleavage, followed by heparin ion-exchange and Superose 6 size-exclusion chromatography. Peak fractions were concentrated by ultrafiltration.

For fluorescent labelling of cohesin the C-terminus of Rad21 was fused to the SNAP tag. SNAP-Rad21 cohesin was purified and labelled with BG-surface Alexa 647 (NEB) as previously described [39]. Labelling was performed after the ion-exchange step, followed by gel-filtration on a Superose 6 10/300 GL (GE Healthcare) column. Labelling efficiency was estimated by recording the absorbance at 280 and 650 nm using a V-550 Spectrophotometer (Jasco). The samples were then aliquoted, frozen in liquid nitrogen and stored at -80°C.

### DNA substrates preparation

λ-phage DNA (48.5 kbp) was terminally labelled with either Digoxigenin or Biotin in a 50 µl reaction containing 1X NEB buffer 2 (NEB), 0.31 mg/ml λ-phage DNA (NEB), 10 units of Klenow Fragment (3’→5’ exo-) of DNA Pol I (NEB), dATP, dGTP (Promega) and either dUTP-Digoxigenin (Jena Bioscience), dCTP (Promega) or dTTP (Promega), dCTP-Biotin (Jena Bioscience). All nucleotides were added to the final concentration of 33 uM each. The mixture was incubated at 37°C for 25 min, followed by inactivation for 20 minutes at 75 °C. Resulting product was cleaned-up from unincorporated nucleotides using Micro Bio-Spin P30 spin column (Bio-Rad).

To prepare DNA with one end labelled with Biotin and the other with Digoxigenin, first a biotinylated oligonucleotide cosL_Phos-bio [5’(Phosphate)-AGGTCGCCGCCC-3’(Biotin)] was ligated to one end of λ-phage DNA. 30 µl reaction contained 0.167 mg/ml λ-phage DNA (NEB), 1.66 µM cosL_Phos-bio oligo and 1x T4 DNA ligase buffer (NEB). The mixture was heated to 70 °C, incubated for 2 minutes and the oligo was annealed by slowly cooling down the mixture to 16 °C for 1 hour. Subsequently, 1mM ATP and 400 Units of T4 DNA ligase were added and the ligation was performed at 16 °C overnight, followed by purification using Micro Bio-Spin P30 column. The resulting product was further labelled with Digoxigenin in a 30 µl reaction containing 3-times diluted ligated DNA, 1X NEB buffer 2 (NEB), 5 units of Klenow Fragment (3’→5’ exo-) of DNA Pol I (NEB), 33 µM of dATP, dGTP, dCTP (Promega) and dUTP-Digoxigenin. Unincorporated nucleotides were removed using Micro Bio-Spin P30 column.

### Microfluidic flow cell fabrication

Flow cells were constructed as previously described [13] with minor modifications. A flow cell containing 5 microfluidic channels was assembled from a silanised coverslip (Marienfeld, High-precision, 24 × 60 mm), a piece of parafilm with precut channels (1 x 20 mm each) and a glass slide with drilled holes.

To create a hydrophobic coating on the surface, coverslips were cleaned and silanised. Cleaning was done by sequential sonication in acetone, 99% ethanol, 0.1 M NaOH for 5 minutes and rinsing with distilled water. Coverslips were further dried using compressed air and plasma cleaned for 5 min. After that, coverslips were submerged into freshly prepared 5% dichlorodimethylsilane (Sigma-Aldrich) dissolved in heptane and incubated for 1 hour. Finally, coverslips were sonicated in chloroform and distilled water (10 minutes each step), blow-dried using compressed air and stored at room temperature in a closed container.

Metal connectors (New England Small Tube Corp) were glued into the holes in the slide using an epoxy glue (Devcon). Slides were cleaned by sequential 5 min sonication in 0.5 % Sodium hypochlorite, 1% Hellmanex and 99% ethanol. After each cleaning step slides were extensively rinsed with distilled water. Slides were further dried at 100°C on a heating plate. After the flow cell assembly, the plastic tubing was attached to the flow cell via metal connectors to form inlets and outlets.

### Flow-assisted DNA loop extrusion assay

A microfluidic channel (approximately 25 µL including the dead volume) was connected to the syringe pump and first rinsed with TBE buffer (Tris 40 mM, NaCl 50mM, EDTA 1mM). Subsequently 50 µL of 0.05 mg/mL of anti-Digoxigenin antibodies (Roche) diluted in TBE were introduced and incubated for at least 20 min and further washed out with 400 µl of TBE. The surface of the flow-cell was passivated against non-specific binding of proteins and DNA by washing in 50 µL of Pluronic (Sigma-Aldrich) 1% solution in TBE and incubation for 5 minutes. The flow-cell was then extensively washed with 2.4 mL of TBE. 50 µL of 1 nM solution of Digoxigenin-labelled DNA in TBE was introduced at 15 µL/min and incubated for 10 minutes, allowing DNA molecules to randomly attach to the surface with DNA ends separated by 1-2 µm. The unbound DNA was removed by washing 60 µL of TBE at 15 µL/min.

Prior to imaging the flow-cell was equilibrated with 50 µl of the imaging buffer (40 mM Tris-HCl pH 7.5, 50 mM NaCl, 2 mM MgCl2, 1 mM ATP, 10 mM DTT, 200 nM Sytox Orange, 1 mg/mL β-Casein, 0.2 mg/ml glucose oxidase, 35 µg/ml catalase and 4.5 mg/ml dextrose) at 15 µl/min. Fission yeast cohesin and cohesin loader were introduced at 1:2 ratio in the imaging buffer at constant flow of 7 µl/min. The flow cell was illuminated with a 561 nm laser through a Nikon SR HP Apo TIRF 100x/1.49 oil immersion objective and a field of view of 167x167 µm containing DNA molecules stained with Sytox Orange was imaged at 1Hz for 10 minutes using a custom-built TIRF microscope, based on Nikon Eclipse Ti2 system. To image DNA the TIRF angle was adjusted to achieve highly inclined and laminated optical sheet (HILO) illumination. Images were recorded using a sCMOS camera (Andor Sona) and saved as TIFF files without compression for further analysis using FIJI ImageJ. All experiments were performed at room temperature.

To visualize cohesin together with DNA, two-colour imaging was performed using Alexa-647 labelled cohesin. Images in two channels were acquired with the OptoSplit (Cairn Research) and two Andor Sona cameras by alternating excitation with 561-nm and 647-nm lasers. For quantification of cohesin stoichiometry, 5nM of fluorescently labelled cohesin and 10nM of Scc2 loader were premixed in the imaging buffer, introduced into the flow cell and incubated for 10 min. In this case, 10mM DTT in the imaging buffer was replaced with 1mM TCEP. To reduce fluorescent background, unbound cohesin was removed by washing the flow-cell with 40 µL of the imaging buffer. After that, cohesin and DNA were imaged at 2Hz with 100 ms exposure time while continuously washing the flowcell with the imaging buffer at 7 µL/min. Edges of the OptoSplit frame were used for the alignment of the two channels for image processing.

### DNA loop extrusion assay without the flow

To visualize DNA loop extrusion without flow, DNA molecules were attached to the surface with ends separated by about 5-6 µm. This allowed enough optical resolution to resolve looped and non-looped regions while keeping DNA tension at minimum. A microfluidic channel was first rinsed with 100 µL of TBE buffer. 40 µL of Avidin-DN (Vector Labs) diluted to 0.1 mg/mL in TBE buffer were introduced into the flowcell and incubated for at least 20 minutes and further washed with 400 µl of TBE. The surface of the flow-cell was passivated by incubating 50 µL of Pluronic 1% solution in TBE for 5 minutes and then extensively washed with 2.4 mL of TBE. Biotinylated DNA diluted to 10pM in TBE was loaded at the constant rate of 4 µL/min for 20 minutes. After that, the channel was immediately washed with 60 µL of TBE at 4 µL/min.

Prior to imaging the flow-cell was equilibrated with 50 µl of the imaging buffer. 40 µL of cohesin and cohesin loader (1:2 ratio in the imaging buffer) were introduced into the channel at 20 µl/min. After that the flow was stop and images were collected at 2 Hz with 100ms exposure time. One field of view (167x167 um) was imaged for 240 seconds, after which the field of view was changed and imaging repeated.

### Optical tweezers assay

Optical trapping experiments were performed on the commercial optical tweezers microscope (Nanotracker 2, JPK) upgraded with a custom-built TIRF setup. The flow-cell was made with longer channels (1 x 4 mm) to accommodate for the condenser lens during force measurement. The flow cell was prepared following the protocol of the flow assisted LE assay using double-labelled (Biotin-Digoxigenin) DNA. After DNA immobilisation, 40 µL of 0.86 um Streptavidin-coated beads (Spherotech) diluted 10 times in the TBE buffer containing 1 mg/mL β-Casein were introduced into the flow cell at 5 µL/min and incubated for at least 30 minutes. Unbound beads were washed out at 5 µL/min with 100 µL of TBE buffer containing 1 mg/mL β-Casein.

The flow-cell was equilibrated with 50 µl of the imaging buffer. A candidate bead was captured using the optical trap. Live fluorescent imaging using HILO illumination was used to verify that the bead was attached to a single DNA molecule. The distance between DNA ends was adjusted to about 2 µm to reproduce conditions of the flow assisted LE assay. The laser power was set to reach the trap stiffness of about 0.002 pN/nm. Cohesin and cohesin loader were diluted to 15 and 16 nM respectively and introduced into the flowcell at 7 µL/min while imaging at 1Hz. Once a DNA loop was formed, the flow was stopped and the distance between the DNA ends was gradually increased by moving a high-precision nanostage at 200 nm/s while simultaneously registering the force at 2kHz and recording images of the DNA at 1-3 Hz. Alternatively, a force-clamp of 0.05 - 0.1 pN was applied and the position of the nanostage was recorded at 400 Hz.

### Data analysis

Fluorescence image series were processed in Fiji/ImageJ (2.16.01/1.54p). In loop extrusion assays, DNA loop extrusion frequency was calculated as the number of double-tethered DNA molecules containing visible loops divided by the total number of double-tethered molecules. Single-tethered molecules and overlapping DNA molecules in which DNA looping determination was obstructed were excluded from analysis.

Loop extrusion analysis in the presence of flow was done as previously described (Higashi et al, 2021). The DNA length prior to loop extrusion was measured in pixels, averaged over a 5 s interval, and normalised by 48.5 kb to calculate a pixel-to-kbp coefficient. To extract the ‘top’ and ‘bottom’ DNA amount, the length of the DNA between the DNA loop and the top or bottom anchor points respectively was measured frame by frame and recalculated in kbp. DNA loop size was calculated by subtracting a sum of ‘top’ and ‘bottom’ DNA from the total length of 48.5 kbp. To estimate the DNA loop extrusion rate, the change of the DNA loop size over time was analysed. A search window of at least 5 seconds long was selected in which the slope of the best linear fit was taken as the loop extrusion rate.

For loop extrusion analysis without the flow, the images were first processed in Fiji/Image. The background was corrected using Subtract background function (50 pixels), followed by Median filter (4 pixels). Subsequently, individual DNA molecules were selected by drawing a line across both DNA ends covering both DNA and a few pixels of the background around it. Kymographs were generated by averaging fluorescence intensity across 12 pixels with the centre on the DNA-marking line using KymoResliceWide plugin (https://doi.org/10.5281/zenodo.4281086). Kymographs were saved as TIFF files and further processed using a custom-made script in Jupyter notebook (Python).

Kymographs were processed line by line, where each line represented a DNA intensity profile at a given time point. Each DNA profile was additionally background corrected by calculating a median filer (3 pixels) and subtracting its minimum value from the original DNA profile. The position of the DNA loop at each time point was determined by finding the maximum in each line of the kymograph. DNA loop size was calculated as a sum of a 13-pixels segment centered on the DNA loop position (loop segment). DNA amount in the ‘top’ and ‘bottom’ segments was calculated by summing pixel values above and below the loop segment respectively. In the absence of flow the DNA loop overlaps the with non-looped DNA. To account for this, the average intensity per pixel in the ‘top’ and ‘bottom’ segments was calculated and multiplied by the size of the loop segment (13 pixels). This value was subtracted from the DNA loop intensity, divided by 2 and each half was added to the ‘top’ and ‘bottom’ DNA intensities. Finally, the total sum of the pixel intensities across the DNA profile was used to recalculate the intensity into kilobasepairs, assuming that total intensity corresponds to 48.5 kbp.

For two-colour experiments, cohesin fluorescence intensity at loop bases was analyzed by photobleaching Alexa-647 fluorophore to estimate the number of cohesin complexes associated with each loop. A kymograph was generated using KymoResliceWide plugin by drawing a 6-pixels wide line centered at the base of the loop. Cohesin signal was extracted from the kymograph by summing 6 pixels around the maximum value in each line of the kymograph. The number of fluorophores was counted as a number of bleaching steps in each trace. In case of no bleaching occurred, the number of fluorophores was determined by comparing the intensity value to a nearby molecule immobilized on the surface that showed single photobleaching step.

For optical tweezers data, bead displacement, DNA tension, and fluorescence time series were analyzed jointly to identify loop formation, force-clamp intervals, and DNA release events from the two flanking arms. Amount of the ‘top’ and ‘bottom’ DNA segments upon application of force was extracted as described for the flow-assisted LE assay. A mutual exclusivity index was computed for each experiment. Each datapoint represented an independent experiment.

### Simulations

We performed coarse-grained molecular dynamics (MD) simulations of a DNA–cohesin system in the NVT ensemble using the LAMMPS software [40]. DNA was represented as a polymer of 1200 beads, with each bead corresponding to 20 base pairs (bp), giving a total contour length equivalent to ∼8 𝜇𝑚. In the simulations length was measured in units of the bead diameter 𝜎 and energy in units of ε = 𝑘_𝐵_T, DNA excluded-volume interactions were modelled by a purely repulsive Weeks–Chandler–Andersen (WCA) potential, connectivity by finitely extensible nonlinear elastic (FENE) bonds, and bending rigidity by a cosine angle potential chosen to reproduce the physical persistence length of DNA. Interactions between cohesin units and DNA were modelled using an attractive Morse potential, implemented through a custom dynamic-window pair style that restricts binding to a local contour region and thereby enforces one-dimensional sliding along a single DNA strand while suppressing unphysical long-range jumps between distant DNA segments. The dynamics were advanced using a velocity-Verlet scheme with time step Δ𝑡 = 0.005𝜏_𝐿𝐽_ according to a Langevin equation

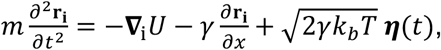

where 𝐫_𝐢_ is the position of bead 𝑖, 𝑈 is the sum of all interaction potentials (as detailed below), and 𝑚 and 𝛾are the mass and the friction coefficient respectively. In the final term 𝜼(𝑡) represents uncorrelated noise of unit variance which provides thermal motion. Here 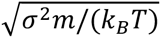 we set 𝑚 = 1 and 𝛾 = 1, and map simulation times to real times via considering the Brownian time scale 𝜏_𝐵𝑟_ = 𝐷/𝜎^2^ where the 3D diffusion constant for a DNA bead 𝐷 can be related to the bead size and fluid viscosity via the Stokes-Einstein equation 𝐷 = 𝑘_𝐵_𝑇/(6𝜋𝜈𝑟). Taking the bead radius as r= 3.4 𝑛𝑚 the viscosity of water 𝜈 ≈ 10^-3^ Pa s leads to a simulation timestep mapping to approximately 3.6 𝑛𝑠.

### Simulation model details

To model excluded-volume, non-bonded DNA beads interact via a purely repulsive WCA interaction given by

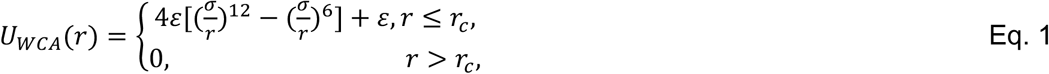

where r is the distance between two beads (implemented in LAMMPS using the lj/cut pair style).

Neighbouring DNA beads are connected by finitely extensible nonlinear elastic (FENE) bonds defined as

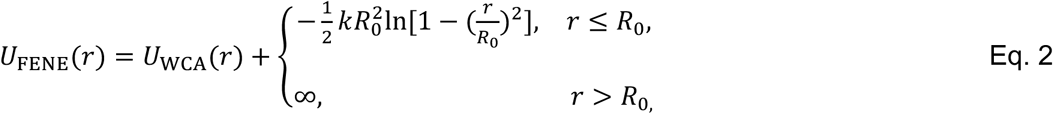

with parameters k = 30ε/𝜎^2^ and 𝑅_0_ = 1.6σ (the LAMMPS “special bonds fene” setting was used to avoid double-counting of excluded volume between bonded neighbors).

DNA bending rigidity was included via a potential

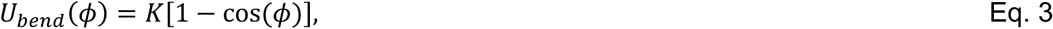

where 𝜙 is the angle made between three consecutive beads in the chain (implemented using the LAMMPS cosine angle style). We set K = 𝑙_𝑝_/b (in units of 𝑘_𝐵_𝑇) to match the physical persistence length of DNA. With 𝑙_𝑝_ ≃ 50 nm (with a discretization of 20 bp per bead, then b = 20 × 0.34 nm = 6.8 nm so K ≃ 50/6.8 ≃ 7.35𝑘_𝐵_𝑇).

### Cohesin/DNA interaction

Cohesin was modelled as two units (motor and anchor) connected by the FENE potential. The motor unit in the motoring state was attached to a specific DNA bead by an explicit harmonic bond, with potential

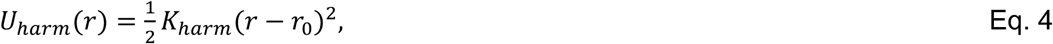

where 𝐾_ℎ𝑎𝑟𝑚_ = 10𝑘_𝐵_𝑇 and 𝑟_0_ = 0.35 𝜎. Directed stepping in the motor state was implemented by deleting the existing harmonic bond and creating a new harmonic bond to a neighboring DNA bead along the contour, thereby producing unidirectional translocation of the motor unit. This was done at regular time intervals such that the motor moved at constant velocity.

In the diffusive state, the harmonic bond was removed and replaced by an attractive Morse interaction between the cohesin unit and DNA, given by

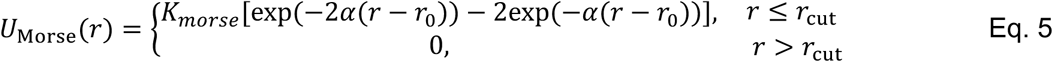

We used 𝐾_𝑚𝑜𝑟𝑠𝑒_ = 14.5 𝛼 = 4.5 𝜎^-1^, 𝑟_0_ = 0.3𝜎, and 𝑟_cut_ = 0.9𝜎. The same interaction potential was used to model diffusive interaction between the anchor and DNA but with variable 𝐾_𝑚𝑜𝑟𝑠𝑒_. This state-dependent interaction scheme was introduced to capture the mechanism suggested by our experimental data and simulations, in which loop extrusion is governed by a motoring unit that can stochastically switch into a diffusive state. The second DNA-binding interface behaves as a diffusive anchor, interacting with the DNA via the same Morse potential as above.

Two cohesin units were connected by a weak FENE tether (as Eq. 2, but with 𝑘 = 0.2𝜀/𝜎^2^ and 𝑅_0_ = 10 𝜎). DNA–cohesin steric interactions were modelled as weak short-range repulsions using the WCA potential (but with an energy 0.5𝑘_𝐵_𝑇, exclusion size 0.1𝜎 and cut off 0.1𝜎) to prevent overlap without introducing strong steric trapping.

To enforce one-dimensional movement of cohesin units in diffusive mode along the DNA, we implemented a custom LAMMPS pair style, morse/dynamic_window. This interaction uses the standard Morse functional form above, but restricts which DNA beads are eligible to exert attractive force on a specified cohesin unit. For a given constrained unit, let 𝑠(𝑡) denote the DNA bead tag identified as the local contour center at the previous timestep. At timestep 𝑡 + Δ𝑡, the Morse attraction is applied only to DNA beads whose contour index 𝑠 satisfies|𝑠 − 𝑠(𝑡)| ≤ 𝑤, where 𝑤 is the contour-window half-width. In all simulations we used 𝑤 = 10 DNA beads. DNA beads outside this contour window exerted no Morse attraction on the corresponding diffusive unit. The window center was updated at every timestep by selecting, among DNA beads within the allowed window, the bead with the smallest Euclidean distance to the cohesin unit and assigning it as the updated center 𝑠(𝑡 + Δ𝑡). At initialization, 𝑠(0) was set to the globally closest DNA bead. Separate dynamic windows were used for the two cohesin units, allowing each to retain its own local contour memory during diffusion.

### Initial DNA conformation and equilibration

For both flow and no-flow simulations, the system was initialized with a DNA polymer consisting of 1200 beads and equilibrated without cohesin for 4 × 10^7^ timesteps before the start of the extrusion simulations. In all cases, the terminal DNA beads (1 and 1200) were kept fixed by excluding them from time integration and setting their forces to zero. The equilibrated DNA conformations were then saved and used as the starting configurations for the cohesin simulations.

For the flow simulations, DNA was initialized in an inverted U-shaped configuration with approximately uniform nearest-neighbor spacing (∼ 0.96𝜎). The DNA contour spanned 𝑥 ∈ [−100,100], with terminal beads located at (−100,0,0) and (100,0,0). During equilibration, a constant external force was applied along the +𝑧 direction to all mobile DNA beads to mimic stretching by flow. The force was chosen such that DNA was stretched to approximately 70% of its contour length, which was similar to DNA length in the experiments in the presence of flow. This captures the effects of the flow in a simple way but does not provide a detailed description of hydrodynamic interactions.

For the no-flow simulations, DNA was initialized in a single-period sinusoidal conformation generated using a custom Python script with approximately uniform nearest-neighbor spacing (∼ 0.96𝜎). In this case, the DNA contour spanned 𝑥 ∈ [−250,250], with terminal beads located at (−250,0,0)and (250,0,0). No external driving force was applied during equilibration.

The difference between simulated and experimental geometry lies in the fact that, in simulations, DNA can theoretically move freely in three-dimensional space, except for the fixed beads on the string. For example, surfaces do not exist in simulations. Although the presence of surfaces alters the system’s free energy (because it restricts certain DNA conformations) and in real experiments, DNA forms an angle with the surface, so the direction of forces should not be entirely along the z-axis, and the forces on each particle cannot be guaranteed to be uniform in experiments, we expect this to have a small impact on our results and will not change our conclusions.

### Simulation protocol and data analysis

As initial condition for all simulations we attached motor and anchor units of cohesin separated by ∼500 bp, which corresponds to the length of DNA with maximum probability of looping spontaneously [41]. The simulations proceeded until cohesin motor reached the end of the DNA. Our Morse potential was chosen such that cohesin experienced small barriers when moving between adjacent DNA beads, which determined its diffusion along the DNA, and much larger barrier for complete dissociation from DNA. Because of this, in some simulations cohesin detached before reaching the end of the DNA when random thermal fluctuations were sufficient to overcome the larger barrier for full dissociation. For analysis, we disregarded all simulations, in which cohesin travelled less than 10% of the total distance. Across all simulations, average number of iterations until cohesin detachment was 1.5 × 10^7^ with 1.8 × 10^7^ required for cohesin to reach the end of the DNA. Thus, in most simulations cohesin completed more than 80% of its movement.

For simulations in which motoring unit was assumed to switch from motoring to diffusive states, we switched motoring to diffusive state after 5.2 × 10^6^ steps. This results in maximum six switches on average before the motoring unit reached the DNA tether point.

To quantify simulated outcomes as symmetric loop extrusion, asymmetric loop extrusion, or translocation we defined a directionality parameter 𝜉 as ratio of the distance travelled by a motor unit towards its nearest DNA tether divided by the direction travelled by the anchor unit towards the opposite DNA tether, quantified at the end of each simulation. Thus, 𝜉=1 corresponds to ideally symmetric loop extrusion when motor and anchor travelled exactly same distance towards the respective DNA tether points, 𝜉=0 corresponds to asymmetric loop extrusion when anchor did not move (travelled zero distance) and 𝜉=-1 corresponds to translocation when motor and anchor travelled the same distance in the same direction (one towards and one away from the corresponding DNA tether points). We then classified all simulated trajectories by imposing a 0.45 value threshold such that all trajectories with 𝜉 > 0.45 we classified as symmetric loop extrusion events, trajectories with −0.45 < 𝜉 < 0.45 as asymmetric events, and trajectories with 𝜉 < −0.45 as translocation events. The value of the threshold (0.45) was chosen such that there was best possible match between simulated and experimental loop extrusion quantified as symmetric.

### Anchor diffusion parameter

To vary the diffusion parameter of anchor cohesin units we changed values of the interaction energy between the anchor and DNA using coefficient 𝐾_𝑚𝑜𝑟𝑠𝑒_ in Eq. 5. The five sets of simulations presented in Fig. 3 correspond to 𝐾_𝑚𝑜𝑟𝑠𝑒_ values of 14.5, 17.5, 20.5, 23.5, and 24.5 𝑘_𝐵_𝑇. Smaller values of 𝐾_𝑚𝑜𝑟𝑠𝑒_ resulted in frequent detachment of cohesin from DNA before reaching the end. At higher values, loop extrusion proceeded asymmetrically in all cases.

To validate the effective diffusion associated with each 𝐾_morse_, we analyzed trajectories of cohesin in the diffusive state on DNA in the absence of flow. For each independent simulation trajectory, the position of the diffusing cohesin unit was represented by the nearest DNA bead index 𝑥(𝑡), obtained from the DNA-contact coordinate. This contour coordinate was converted to physical distance using 6.8 nm per DNA bead.

For each trajectory, the squared displacement was calculated relative to the first analyzed frame, Δ𝑥^2^(𝑡) = [𝑥(𝑡) − 𝑥(𝑡_0_)]^2^, where 𝑡_0_ denotes the first sampled time point of that trajectory. The mean-squared displacement (MSD) was then obtained by averaging Δ𝑥^2^(𝑡)across independent simulation trajectories at each sampled time point. Error bars represent the standard error of the mean across trajectories.

Simulation time was converted to physical time using 1timestep = 3.6 × 10^-R^s. Diffusion coefficients were extracted by linear fitting of MSD versus time using the one-dimensional diffusion relation MSD(𝑡) = 2𝐷𝑡. For 𝐾_morse_ = 14.5 𝑘_𝐵_𝑇 and 17.5 𝑘_𝐵_𝑇, the fit was restricted to the initial approximately linear regime (0–600 ms and 0–280 ms, respectively), whereas for 𝐾_morse_ = 20.5, 23.5, and 24.5 𝑘_𝐵_𝑇, the full sampled time range was used. The resulting 1D diffusion coefficients were used in Fig. 3c–e.

### Computational analysis and hardware

The simulations and analysis were run on a local workstation equipped with an Intel Core i9-14900K CPU (24 cores, 32 threads), 128 GB DDR5 RAM, and a 960 GB SSD, running Ubuntu 20.04.5 LTS (GNU/Linux 5.4.0-216-generic, x86_64), using lammps-22Jul2025.

## Author contributions

GP performed microscopy and optical trapping experiments, RS and CB performed simulations; TH and FU designed, expressed and purified proteins used in the study; OD performed protein expression, purification and quality control; GP and MM designed the study and wrote the manuscript.

## Acknowledgments

This study was supported by the Francis Crick Institute, which received funding from the UK Medical Research Council, Cancer Research UK and the Wellcome Trust through award FC001750 to MM.

## Code availability

LAMMPS code used for the simulations is freely available in the GitHub repository https://github.com/FrancisCrickInstitute/DiffusiveCohesin/. Image processing code for kymographs is available at https://github.com/PobegalovG/Pobegalov-et-al.cohesin-LE-kymograph-analysis/.

## Supplementary Video Legends

**Video S1.** Example of cohesin DNA loop extrusion in the presence of flow. Scale bar – 2 microns.

**Video S2.** Example of cohesin DNA loop extrusion in the absence of flow. Scale bar – 2 microns.

**Video S3.** Example of the simulation for the model with continuous motor (continuously moving towards one of DNA tether points) and diffusive anchor resulting in asymmetric loop extrusion. Diffusion coefficient of the anchor is 0.3 𝜇m^2^/s.

**Video S4.** Example of the simulation for the model with continuous motor and diffusive anchor resulting in translocation. Diffusion coefficient of the anchor is 1 𝜇m^2^/s.

**Video S5.** Example of the simulation for the model with continuous motor and diffusive anchor resulting in near symmetric loop extrusion. Diffusion coefficient of the anchor is 1 𝜇m^2^/s.

**Video S6.** Example of the simulation for the model with stochastically state-switching motor and diffusive anchor resulting in symmetric loop extrusion. Diffusion coefficient of the anchor is 0.3 𝜇m^2^/s.

**Video S7.** Example of pulling on cohesin-mediated DNA loop using optical tweezers.

**Video S8.** Example of DNA loop extrusion by Scc3-Nhp6A fusion cohesin.

---

**Supplementary Figure 1.**
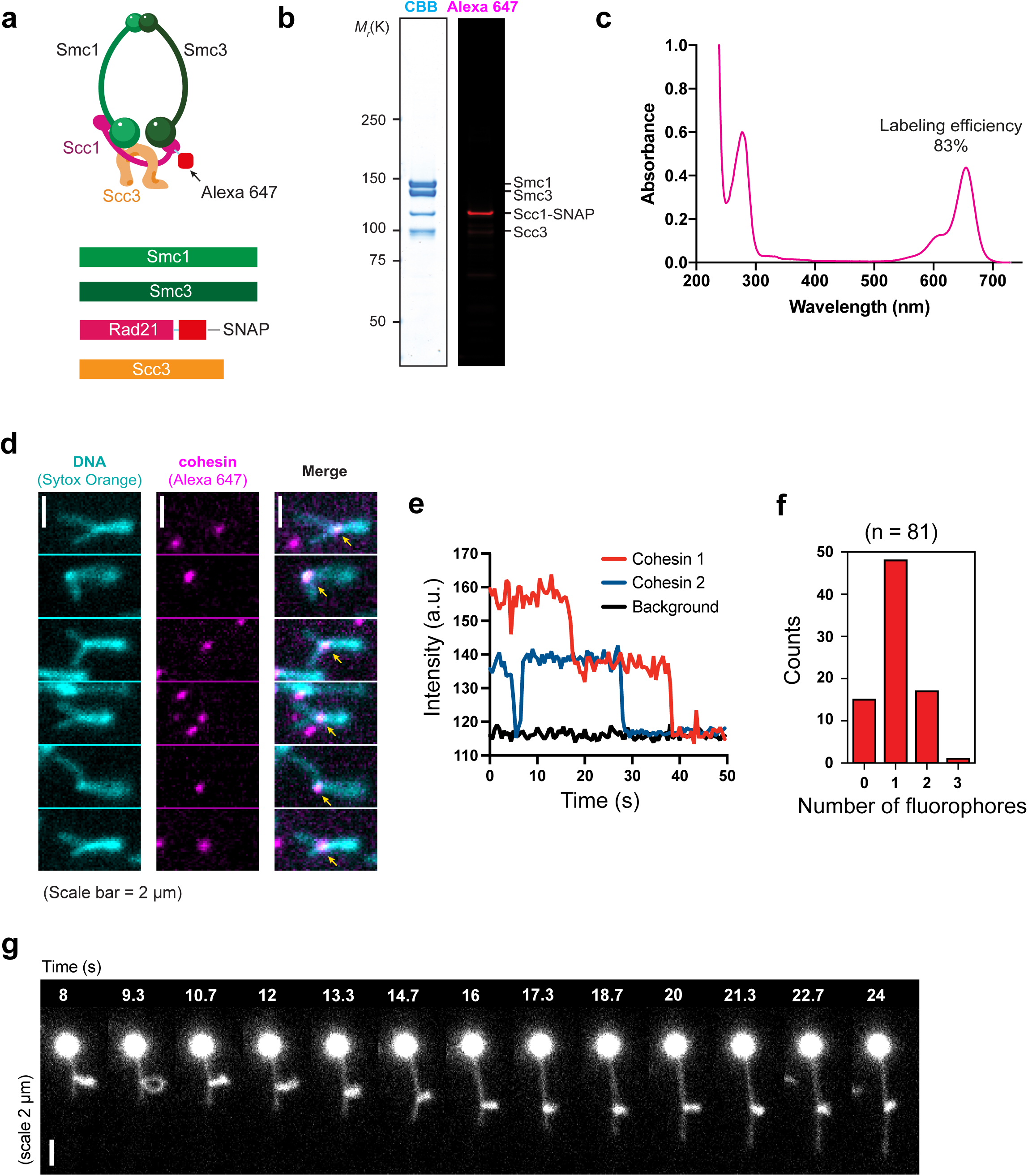
Labelled cohesin preparation and quantification. (a) Schematics of the SNAP-tagged cohesin complex used for Alexa-647 labelling; (b) Left, SDS-PAGE gel stained with Coomassie Blue showing the purified subunits of the cohesin. Right, in-gel Alexa-647 fluorescence of the Scc1-SNAP subunit, confirming successful Alexa-647 attachment; (c) Example absorbance spectrum of 2 µM Alexa647-cohesin. The labelling efficiency was ∼86%, assuming the molar extinction coefficient for the Alexa-647 dye of 270,000 M-1cm-1; (d) Representative images showing single cohesin molecule at the base of the DNA loop. Yellow arrows point to cohesin; (e) Two example photobleaching traces showing bleaching of a single (blue) or double (red) cohesin at the base of the loop; (f) Histogram showing distribution of the cohesin photobleaching events; (g) Representative fluorescence time series of pulling on a DNA loop with the side flow to visualize the loop. Scale bar, 2 µm.

**Supplementary Figure 2.**
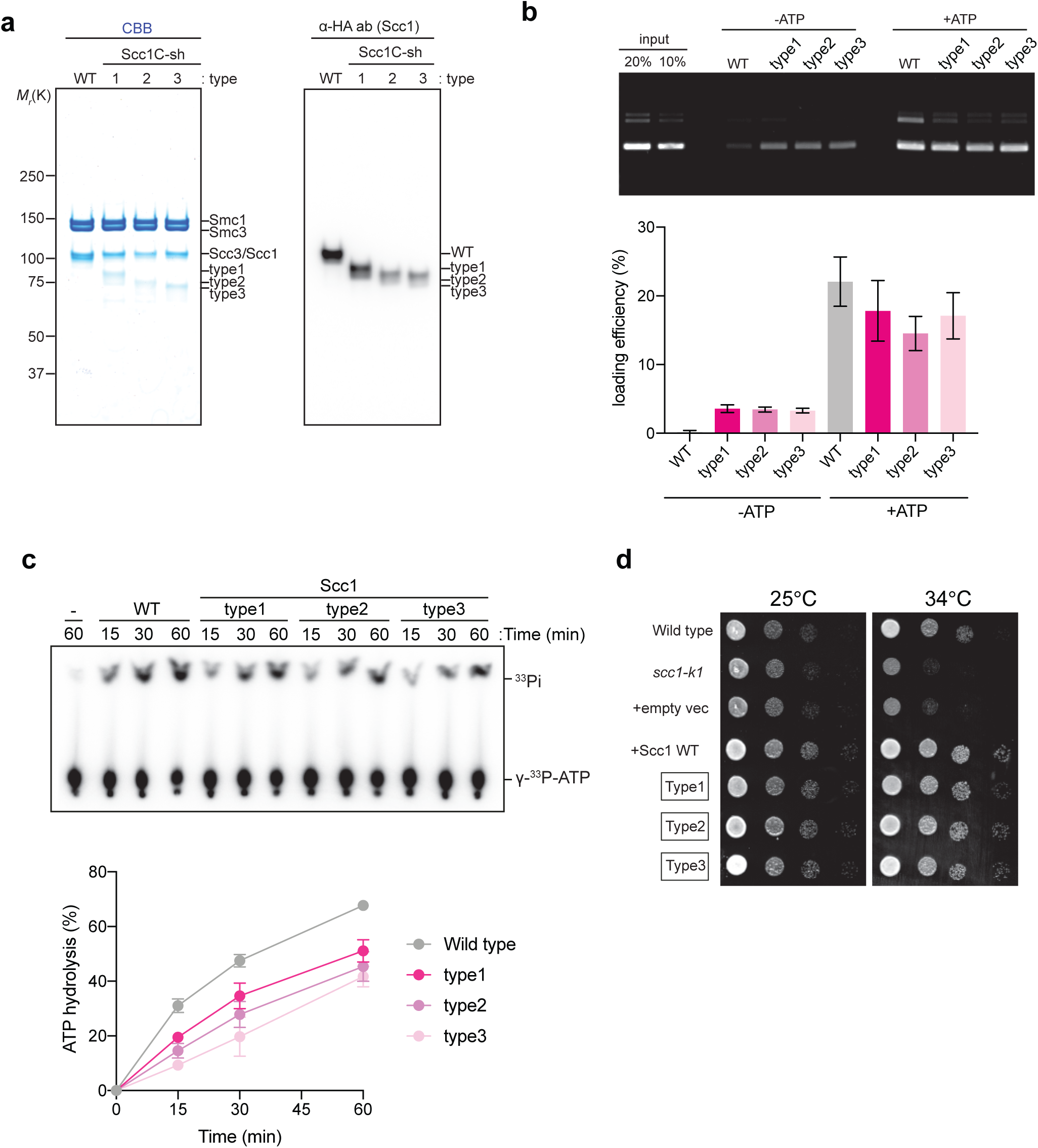
Characterization of Cohesin Scc1 truncation mutants. (a) Purified fractions analysed by SDS-PAGE followed by Commiassie blue staining or Western blotting against the HA epitope tag that was fused to Scc1. Type 1, 2 and 3 mutants correspond to truncations with the expected lengths of Scc1 C-terminus of 19, 9.5 and 4.5 nm correspondingly; (b) Cohesin complexes harbouring truncated Scc1 topologically entrap DNA similarly to the wild type complex. A topological DNA loading assay was performed in the absence or presence of ATP, and the fraction of input DNA bound by cohesin in a salt resistant manner quantified. Three biological repeats of the experiment were performed, the mean and standard deviations are shown; (c) ATP hydrolysis rate measured for the indicated cohesin variants; (d) Cohesin complexes harbouring truncated Scc1 support cell viability. Serial dilutions of strains of the indicated genotypes were spotted on YES agar plates and incubated at the indicated temperatures.

**Supplementary Figure 3.**
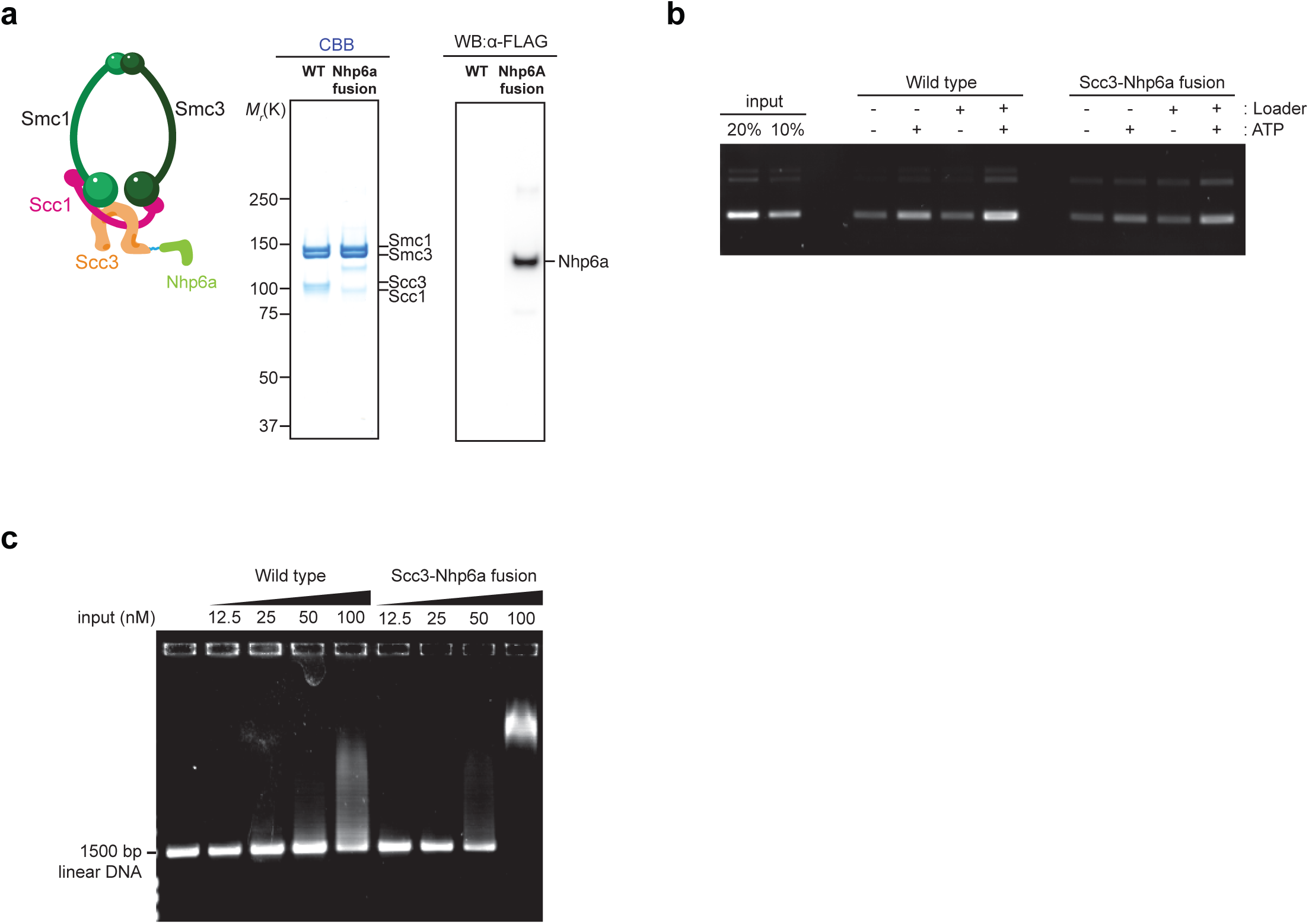
Construction and characterisation of cohesin with Nhp6A fused to the Scc3 subunit. (a) Schematic depicting the design of the fusion. Cohesin complexes containing Scc1 fused to Nhp6A was expressed and purified and the final purified complex analysed by SDS-PAGE followed by Commiassie blue staining or Western blotting against the FLAG peptide that was included in the Nhp6A fusion; (b) Cohesin complex harbouring Scc3-Nhp6A fusion topologically entrap DNA similarly to the wild type complex. A topological DNA loading assay was performed in the absence or presence of the indicated components; (c) Increased DNA affinity of cohesin harbouring the Scc3-Nhp6A fusion. An electrophoretic gel mobility shift assay was performed using the indicated concentrations of wild type or fusion cohesin complexes

**Supplementary Figure 4.**
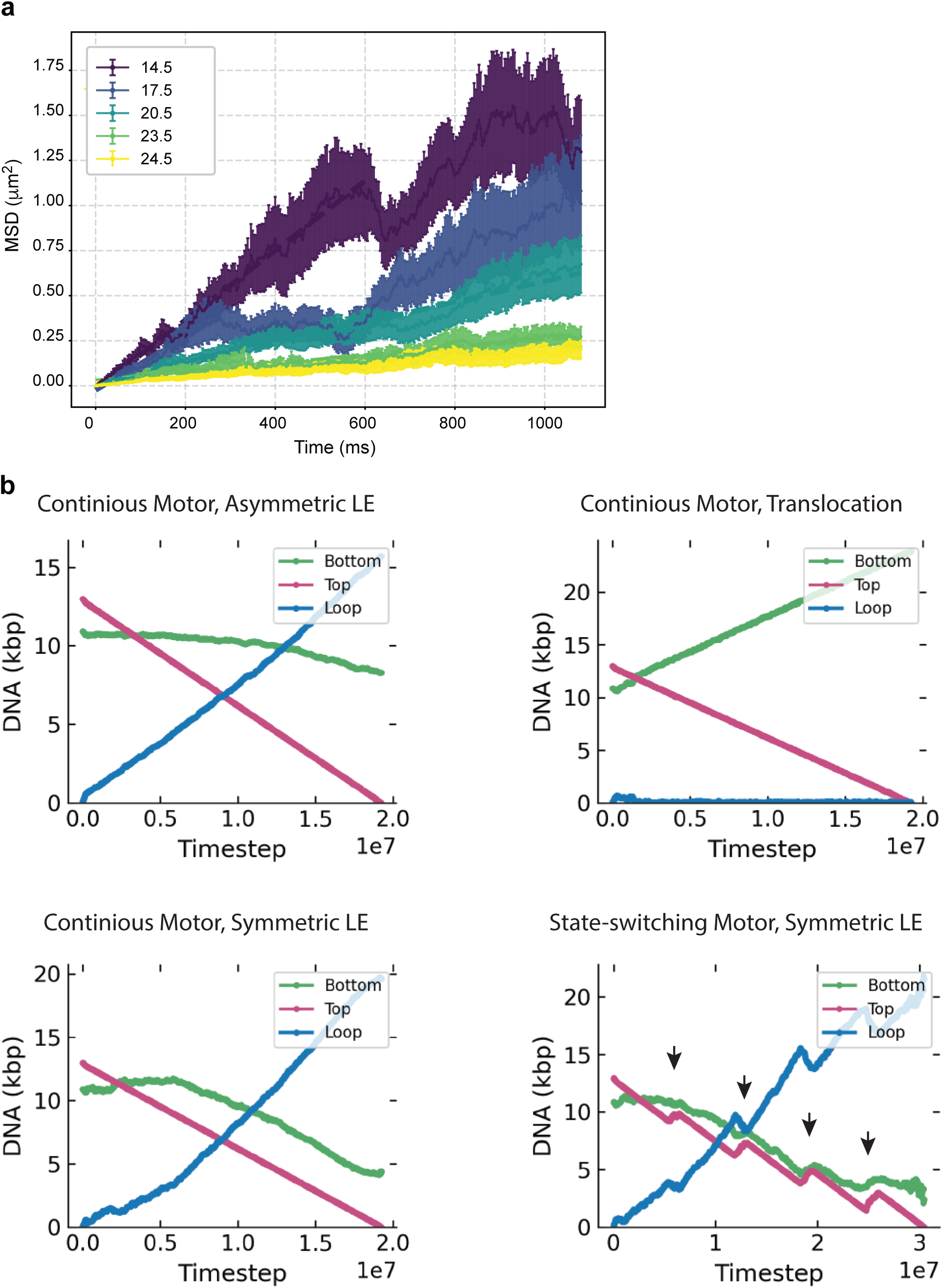
Quantification of the diffusion coefficients and examples of simulations. (a) MSD plots for cohesin diffusing on DNA without the flow used to extract diffusion coefficients for Fig. 2. Colours correspond to different strength of the anchor/DNA interaction. Shown values correspond to values of parameter *K_morse_*. Diffusion values were obtained by fitting initial 200 ms of these plots; (b) Time traces showing the lengths of the top and bottom DNA arms as well as the size of the loop corresponding to simulations in videos S3, S4, S5 and S6. Events labelled with arrows correspond to switching of the motor between motoring and diffusing states.

